# Classification of RNA-Seq Data via Bagging Support Vector Machines

**DOI:** 10.1101/007526

**Authors:** Gokmen Zararsiz, Dincer Goksuluk, Selcuk Korkmaz, Vahap Eldem, Izzet Parug Duru, Turgay Unver, Ahmet Ozturk

**Author notes:** Corresponding author Email addresses: GZ DG SK VE IPD TU AO.

## Abstract

**Background:** RNA sequencing (RNA-Seq) is a powerful technique for transcriptome profiling of the organisms that uses the capabilities of next-generation sequencing (NGS) technologies. Recent advances in NGS let to measure the expression levels of tens to thousands of transcripts simultaneously. Using such information, developing expression-based classification algorithms is an emerging powerful method for diagnosis, disease classification and monitoring at molecular level, as well as providing potential markers of disease. Here, we present the bagging support vector machines (bagSVM), a machine learning approach and bagged ensembles of support vector machines (SVM), for classification of RNA-Seq data. The bagSVM basically uses bootstrap technique and trains each single SVM separately; next it combines the results of each SVM model using majority-voting technique.

**Results:** We demonstrate the performance of the bagSVM on simulated and real datasets. Simulated datasets are generated from negative binomial distribution under different scenarios and real datasets are obtained from publicly available resources. A deseq normalization and variance stabilizing transformation (vst) were applied to all datasets. We compared the results with several classifiers including Poisson linear discriminant analysis (PLDA), single SVM, classification and regression trees (CART), and random forests (RF). In slightly overdispersed data, all methods, except CART algorithm, performed well. Performance of PLDA seemed to be best and RF as second best for very slightly and substantially overdispersed datasets. While data become more spread, bagSVM turned out to be the best classifier. In overall results, bagSVM and PLDA had the highest accuracies.

**Conclusions:** According to our results, bagSVM algorithm after vst transformation can be a good choice of classifier for RNA-Seq datasets mostly for overdispersed ones. Thus, we recommend researchers to use bagSVM algorithm for the purpose of classification of RNA-Seq data. PLDA algorithm should be a method of choice for slight and moderately overdispersed datasets. An R/BIOCONDUCTOR package MLSeq with a vignette is freely available at http://www.bioconductor.org/packages/2.14/bioc/html/MLSeq.html

## Background

With the advent of high-throughput NGS technologies, transcriptome sequencing (RNA-Seq) has become one of the central experimental approaches for generating a comprehensive catalog of protein-coding genes and non-coding RNAs and examining the transcriptional activity of genomes. Furthermore, RNA-Seq has already proved itself to be a promising tool with a remarkably diverse range of applications; (i) discovering novel transcripts, (ii) detection and quantification of spliced isoforms, (iii) fusion detection, (iv) reveal sequence variations (e.g, SNPs, indels) [1]. Additionally, beyond these general applications, RNA-Seq holds great promise for gene expression-based classification to identify the significant transcripts, distinguish biological samples and predict clinical or other outcomes due to large amounts of data which can be generated in a single run.

Although microarray-based gene expression classification have become very popular during last decades, more recently, RNA-Seq replaced microarrays as the technology of choice in quantifying gene expression due to some advantages as providing less noisy data, detecting novel transcripts and isoforms, and unnecessity of prearranged transcripts of interest [2–5]. Additionally, to measure gene expression, microarray technology provides continuous data, while RNA-Seq technology generates discrete count data, which corresponds to the abundance of mRNA transcripts [6]. Therefore, novel approaches based on discrete probability distributions (e.g. poisson, negative binomial) are urgently required to deal with huge amount of data for expression-based classification purpose. Another choice is to use some transformation approaches (e.g. vst –variance stabilizing transformation- or rlog –regularized logarithmic transformation-) to bring RNA-seq samples hierarchically closer to microarrays and apply known algorithms for classification applications [7–9]. Recently, a few studies were performed to classify the sequencing data. Witten et al. [6] proposed a Poisson linear discriminant analysis (PLDA) classifier, which is similar to diagonal linear discriminant analysis and can be applied to sequencing data. Ghaffari et al. [10] modelled the gene expression levels as a multivariate normal distribution model by a transformation through a Poisson filtering and tested the performance of linear discriminant analysis (LDA) by using three nearest neighbors and radial basis function of support vector machines (SVM) classifiers. Ryvkin et al. [11] developed a random forest (RF) based classification approach to differentiate six different class of non-coding RNAs on the basis of small RNA-Seq data. In addition to these methodologies, Cheng et al. [12] proposed binomial mixture models to classify bisulfite-sequencing data for DNA methylation profiling.

In this study, we describe bagging support vector machines (bagSVM) as a first use of machine learning algorithms for the purpose of RNA-Seq data classification. The bagSVM is an ensemble machine learning approach, which randomly selects training samples with bootstrap technique, trains each single SVM separately and combines the results of each model to improve the accuracy and the reliability of predictions. This method is applied in several studies to improve the classification performance of SVM [13–16].

## Results and Discussion

### Datasets

A comprehensive simulation is made to compare the performances of the classifiers. In addition to the simulated data, two real datasets were also used to illustrate the usefulness of bagSVM algorithm in real life examples.

*Simulated Datasets:* Simulated datasets are generated under 144 different scenarios using a negative binomial model as follows:

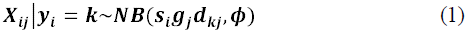

where, *s_i_* is the number of counts per sample, *g_j_* is the number of counts per gene, *d_jk_* is the *j^th^* differentially expressed gene between classes and *φ* is the dispersion parameter. The datasets contain all possible combination of:

- number of biological samples (*n*) changing as 20, 40, 60, 80;
- number of differentially expressed genes (*p*) as 25, 50, 75, 100;
- number of classes (*k*) as 2, 3, 4;
- different dispersion parameters as *φ*=0.01 (very slightly overdispersed), *φ*=0.1 (substantially overdispersed), *φ*=1 (highly overdispersed).

Each gene had a 40% chance of being differentially expressed among classes. *s_i_* and *g_j_* are distributed identically and independently as *s_i_* and *g_j_* respectively.

*Real Datasets:* Two real RNA-Seq datasets are used in this study. A very short data description is listed in Table 1. More details about these datasets can be found in related papers.

**Table 1.** Description of real RNA-Seq datasets used in this study

| Dataset | Description |
| --- | --- |
| *Liver and kidney*<br>[17] | Liver and kidney data measures the gene expression levels of 22,935 genes belonging to human kidney and liver RNA-Seq samples. There were 14 kidney and liver samples (7 replicates in each) and these samples are treated as two distinct classes |
| *Cervical cancer*<br>[18] | Cervical cancer data measures the expressions of 714 miRNA's of human samples. There are 29 tumor and 29 non-tumor cervical samples and these two groups are treated as two separate classes |

### Classification Properties and Results

A DESeq normalization [19] was applied to each dataset to adjust sample specific differences. Variance stabilizing transformation (vst) [19] was performed for each algorithm, except PLDA. For real datasets, differential expression was performed and genes are ranked from most significant to less with increasing number of genes in steps of 20 up to 200 genes. Number of bootstrap samples was set as 100 for the bagSVM models since small changes were observed over 100 bootstraps. Radial basis function was used for the SVM models as kernel and number of trees was set to 100 for the RF models. Other model parameters were optimized using the trainControl function of R package caret [20]. To validate each model, 5-fold cross-validation was used, repeated 10 times and accuracy rates were calculated to evaluate the performance of each model. Simulation results are demonstrated in Figure 1.

**Figure 1.**
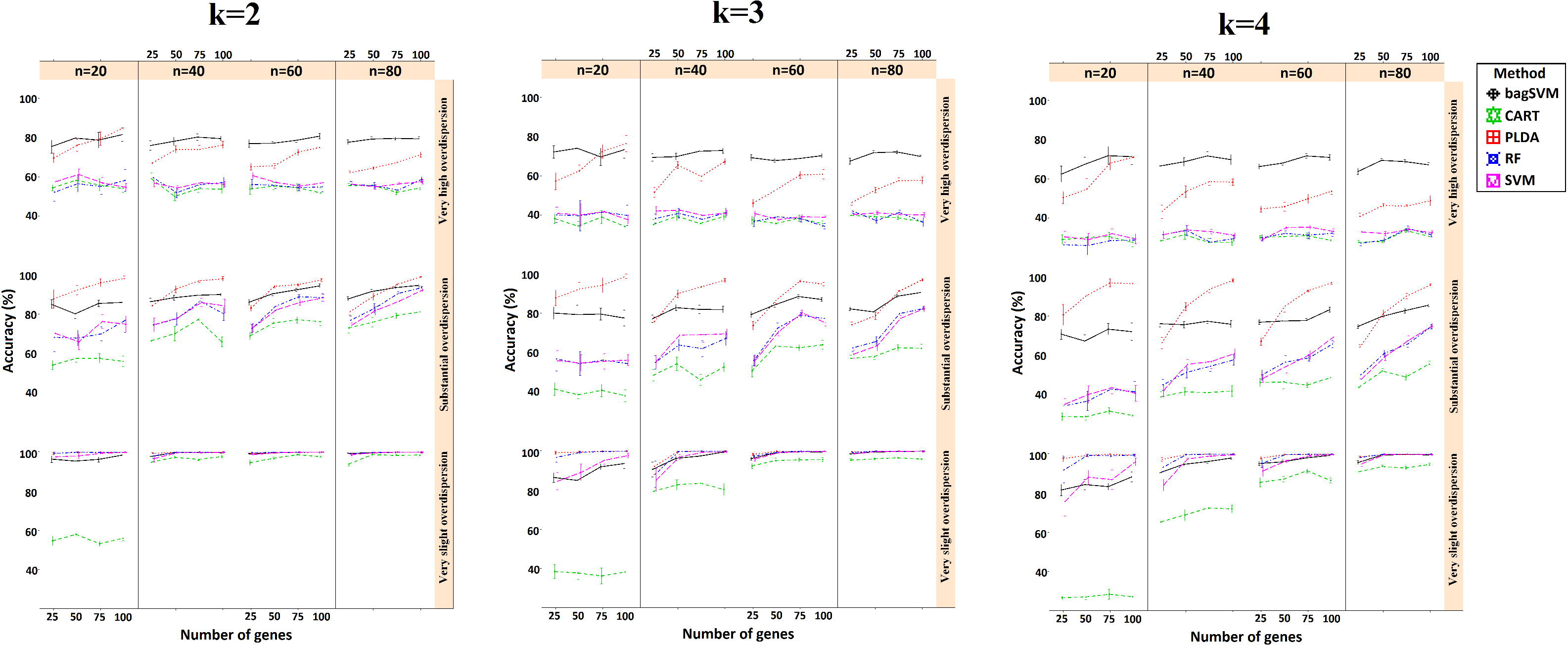
Simulation results with *k*=2, 3 and 4. Figure shows the performance results of classifiers with changing parameters of sample size (*n*), number of genes (*p*) and type of dispersion (*φ*=0.01: very slight, *φ*=0.1: substantial, *φ*=1: very high)

As can be seen from Figure 1, overall accuracies decrease as the number of classes increases. This result is due to the fact that the misclassification probability of an observation may be arised depending on the increase in class number. Besides, the performance of each method was decreasing depending on the increase in overdispersion parameter. In slightly overdispersed data sets, all methods performance was very high, except CART algorithm for small sample sized data. Performance of PLDA seemed to be best and RF as second best for very slightly and substantially overdispersed datasets. The reason may be that PLDA classifies the data using a model based on Poisson distribution. We believe that, extending this algorithm with negative binomial distribution may increase its performance for highly overdispersed datasets. However, it is beyond scope of the present study and we leave this issue as an open question to be addressed in future work.

For substantially overdispersed data, one may choose bagSVM when working with small number of genes, whereas PLDA should be preferred over bagSVM if there are enough number of genes and samples. However, the accuracy results of bagSVM were still high for moderate and slightly overdispersed data (mean accuracies were 82% and 96% respectively) and still can be a preferred classifier in this situation.

While data become more spread, bagSVM turned out to be the best classifier (Figure 1).

Unless we work with technical replicates, RNA-Seq data is overdispersed and for same gene, counts from different biological replicates have variance, which exceeds the mean [21]. This overdispersion can be seen in other research studies [22–26]. Results of our study revealed that overdispersion has a significant effect on classification accuracies and should be taken into account before model building. Moreover, we reach a conclusion that the effect of sample and gene numbers on accuracy rates is largely dependent on the dispersion parameter. In other words increasing number of samples and genes leads to a significant increase in accuracy, unless the data is overdispersed (Figure 1). When data is overdispersed, increasing the number of samples does not change the performance of classifiers. However, increasing number of features significantly increases overall model accuracies in most scenarios.

The results of real data sets is shown in Figure 2. In liver and kidney data, most of the methods performed well except CART algorithm. In cervical cancer data, bagSVM showed the best results and mostly improved the performance of single SVM classifier. Likely in simulation results, bagSVM performed as the best classifier for overdispersed data and all methods performances were very high except CART algorithm for a slightly overdispersed data. The distribution of overdispersion parameter is demonstrated in Figure 3. As seen from the histogram plots, cervical data is a highly overdispersed and liver and kidney data is a lowly overdispersed data. As can be noticed that, results obtained from both real and simulation data sets were consistent with each other (Figure 2). Consequently, we suggest that the bagSVM performed well in both real data sets. It outperformed other algorithms for overdispersed cervical data. Consequently, we conclude that the high performance of classifiers in liver and kidney data may be arised as a result of very low overdispersion as well as small sample size.

**Figure 2.**
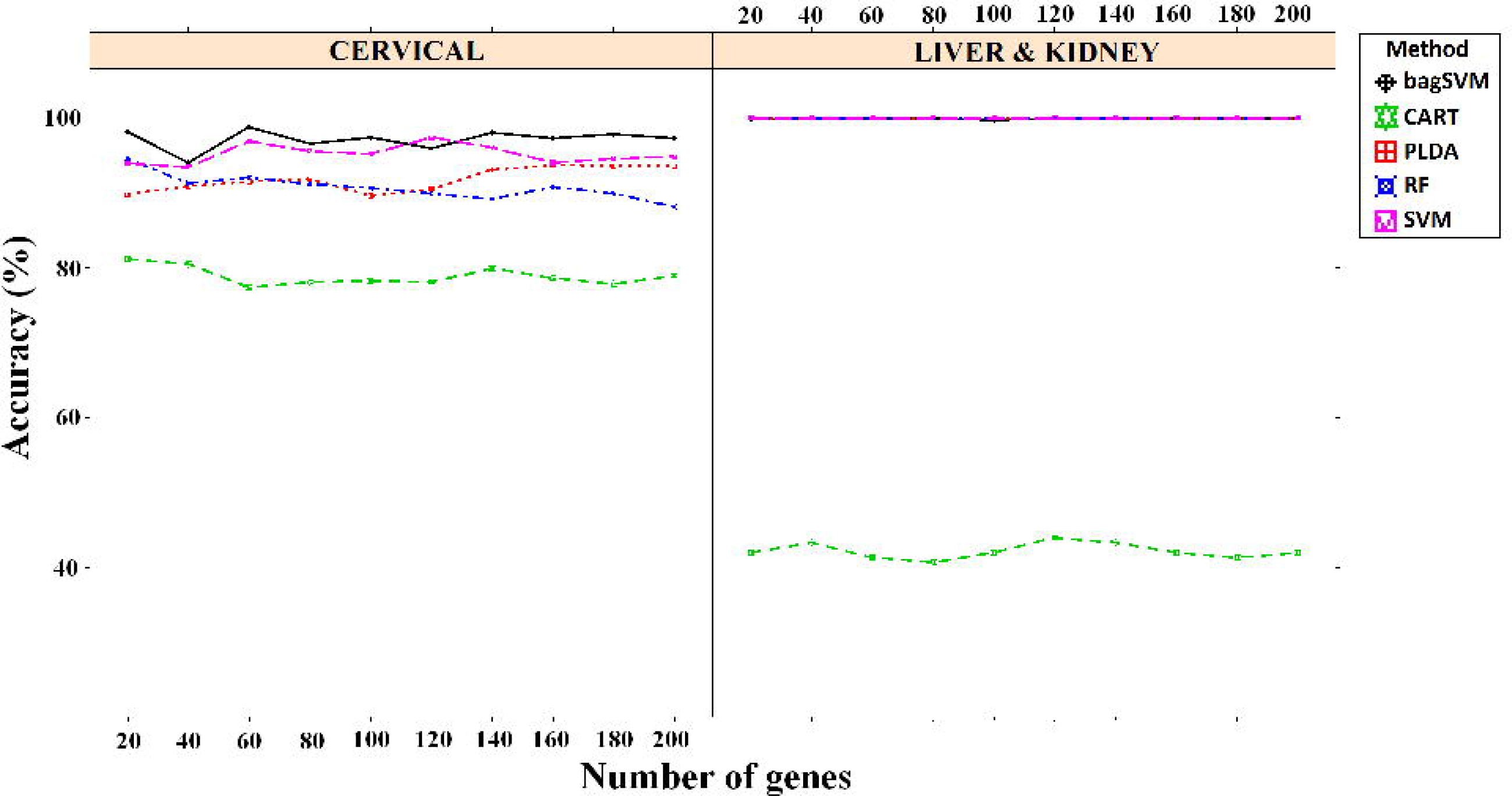
Results obtained from cervical (left panel) and liver & kidney (right panel) real datasets. Figure shows the performance results of classifiers for datasets with changing number of most significant number of genes

**Figure 3.**
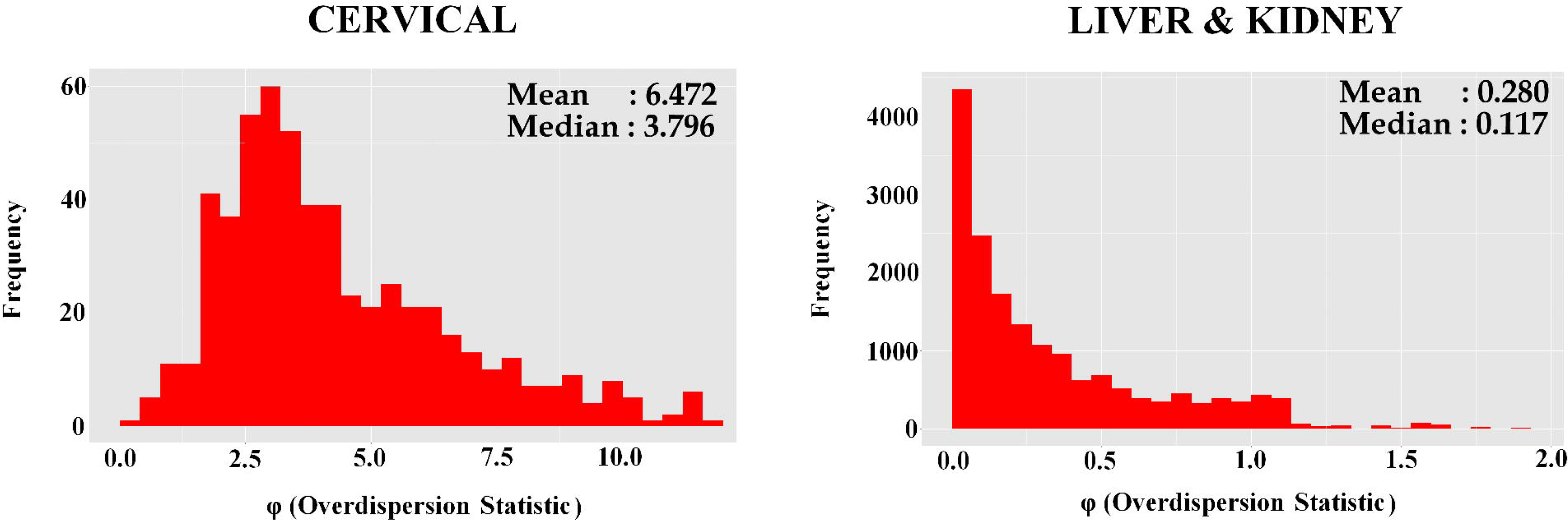
Distribution of overdispersion statistic in cervical and liver & kidney data

To make an overall assessment of the results, we generated error bar plots of classification accuracy results obtained from simulated data sets. In addition, for each scenario, the classifiers were ranked from best to worst based on the classification accuracies. Next, Pihur’s cross-entropy Monte-Carlo rank aggregation approach [27] was applied to get a combined super list to indicating overall performance of each classifier. Finally, overall performances were clustered by hierarchical clustering to see the similarities of classifiers and the results are given in Figure 4. Results revealed that PLDA and bagSVM had the highest accuracies and found to be similar on average, SVM and RF performed moderately similar, while CART performed slightly better than a random chance. A possible explanation for such observation is that bagSVM uses bootstrap technique and trains a model from a dataset which have lower variance. However, it aggregates single models and transform good predictors into nearly optimal ones [28].

**Figure 4.**
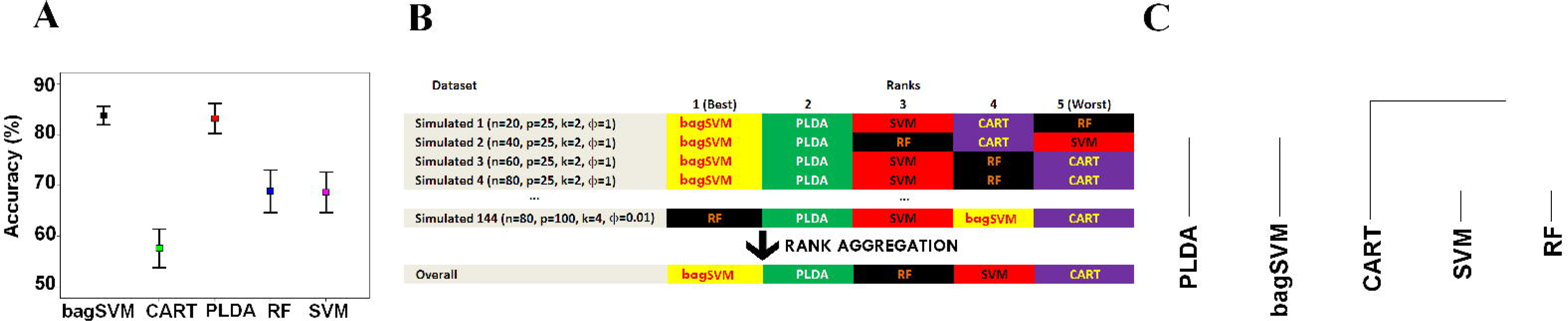
(A) Error bars indicating overall performances of classifiers (B) Pihur’s rank aggregation approach results (C) Hierarchical clustering for euclidean distance and average method to demonstrate overall performance of classifiers

In overall, the performance of SVM and RF were not as high as bagSVM or PLDA, and decrease when the data becomes more spread. It can be said that bagSVM increases the performance of single SVMs when data is barely separable. This result is consistent with the findings of [29–30].

Witten et al. [6] mentioned that normalization strategy has little impact on the classification performance but may be important in differential expression analysis. However, data transformation has a direct effect on classification results, by changing the distribution of data. In this study, we used deseq normalization with variance stabilizing transformation and had well results with bagSVM algorithm. We leave the effect of other transformation approaches (such as rlog [9] and voom [31]) on classification as a topic for further research.

## Conclusions

A considerable amount of evidence collected from genome-wide gene expression studies suggests that the identification and comparison of differentially expressed genes have been a promising approach of cancer classification for diagnosis and prognosis purposes. Although microarray-based gene expression studies through a combination of classification algorithms such as SVM and feature selection techniques have recently been widely used for new biomarkers for cancer diagnosis [32–35], it has its own limitations in terms of novel transcript discovery and abundance estimation with large dynamic range. Thus, one choice is to utilize the power of RNA-Seq techniques in the analysis of transcriptome for diagnostic classification to surpass the limitations of microarray-based experiment. As mentioned in earlier sections, working with less noisy data can enhance the predictive performance of classifiers, and the novel transcripts may be a biomarker in interested disease or phenotypes.

This study is among the first studies for RNA-Seq classification. However, it can be extended to other sequencing studies such as DNA or ChIP-sequencing data. Results from simulated data sets revealed that, bagSVM method performs as the best algorithm when the data is becoming to be overdispersed. When overdispersion is substantial or low, PLDA method becomes an appropriate classifier.

In summary, bagSVM algorithm after vst transformation can be a good choice of classifier for all kinds of RNA-Seq datasets, mostly for overdispersed ones. PLDA algorithm should be a method of choice for slight and moderately overdispersed datasets.

We have developed an R package named MLSeq to implement the algorithms discussed here. This package is publicly available at BIOCONDUCTOR (http://www.bioconductor.org/packages/2.14/bioc/html/MLSeq.html).

## Methods

### Bagging Support Vector Machines

Ensemble methods are learning algorithms that improve the predictive performance of a classifier. An ensemble of classifiers is a collection of multiple classifiers whose individual decisions are combined in some way to classify new data points [36, 37]. It is known that ensembles often represent much better predictive performance than the individual classifiers that make them up [37, 38].

The SVM generally shows a good generalization performance and is easy to learn exact parameters for the global optimum. On the other hand, due to the practical SVM has been carried out using the approximated algorithms in order to reduce the computation complexity, a single SVM classifier may not learn exact parameters for the global optimum [39]. To deal with this issue, several authors proposed to use a bagging ensemble of SVM [29, 38].

BagSVM is a bootstrap ensemble method, which creates individuals for its ensemble by training each SVM classifier (learning algorithm) on a random subset of the training set. For a given data set, multiple SVM classifiers are trained independently through a bootstrap method and they are aggregated via an aggregation technique. To construct the SVM ensemble, K replicated training sets are generated by randomly re-sampling, but with replacement, from the given training set repeatedly. Each sample, ***x_i_***, in the given training set, ***TR***, may appear repeated times, or not at all, in any particular replicate training set. Each replicate training set will be used to train a specific SVM classifier. The general structure of bagSVM is given in Figure 5 and Additional file 2 shows the pseudo-code of the used bagging algorithm.

**Figure 5.**
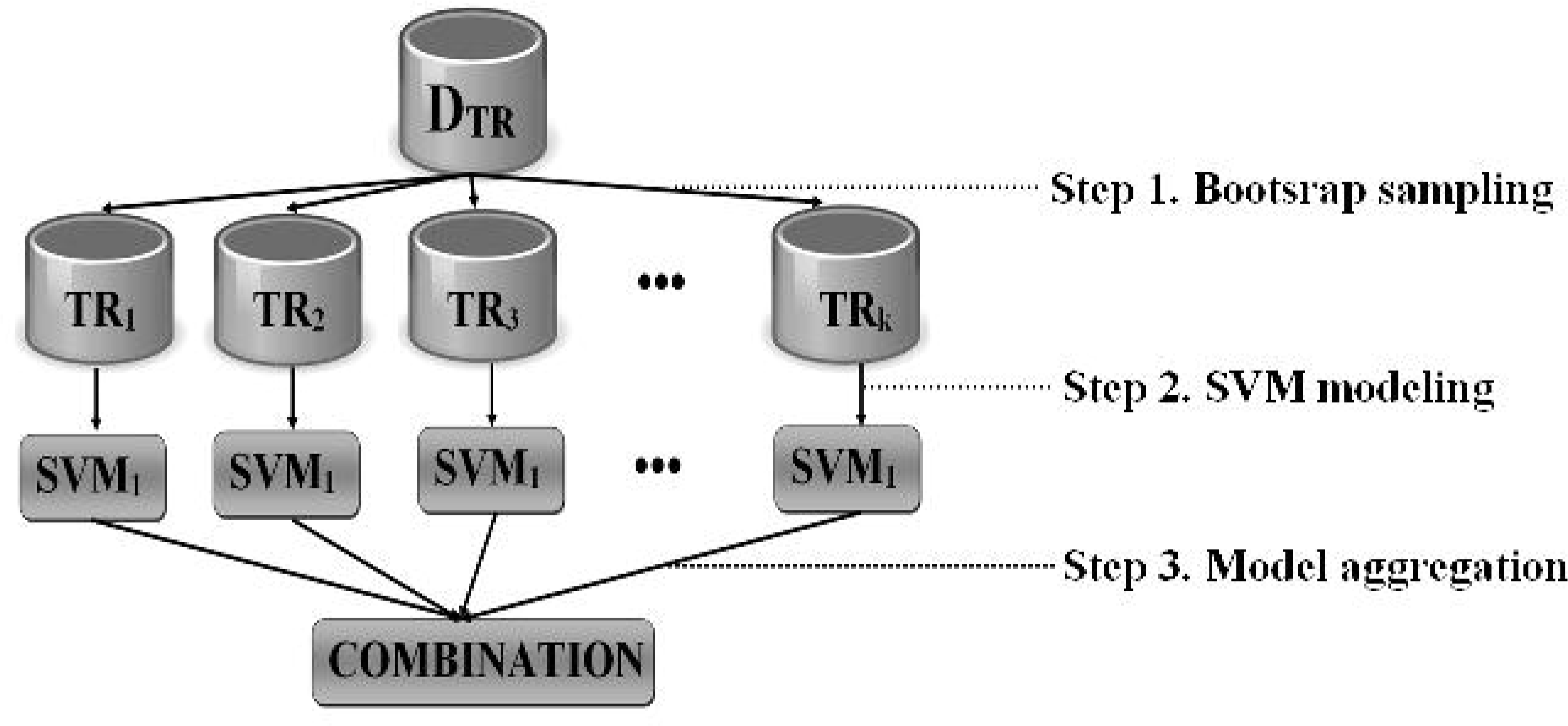
The general structure of bagSVM

### Other Classification Algorithms

*Support vector machines:* SVM is a classification method based on statistical learning theory, which is developed by Vapnik [40] and his colleges, and has taken great attention because of its strong mathematical background, learning capability and good generalization ability. Moreover, SVM is capable of nonlinear classification and deal with high-dimensional data. Thus, it has been applied in many fields such as computational biology, text classification, image segmentation and cancer classification [29, 40].

In linearly separable cases, the decision function that correctly classifies the data points by their true class labels represented by:

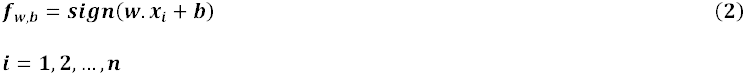

In binary classification, SVM finds an optimal separating hyperplane in the feature space, which maximizes the margin and minimizes the probability of misclassification by choosing ***w*** and ***b*** in equation (2). For the linearly non-separable cases, slack variables {***ξ***_**1**_,…,***ξ_n_***}, which is a penalty introduced by Cortes and Vapnik [41], can be used to allow misclassified data points, where ***ξ*** > 0. In many classification problems, the separation surface is nonlinear. In this case, SVM uses an implicit mapping **Φ** of the input vectors to a high-dimensional space defined by a kernel function (***K***(***x, y***) = **Φ**(***x_i_***)**Φ**(***x_j_***)) and the linear classification then takes place in this high-dimensional space. The most widely used kernel functions are linear: ***K***(***x, y***) = ***x_i_x_j_***, polynomial: ***K***(***x, y***) = (***x_i_x_j_*** + 1)***^d^***, radial basis function: 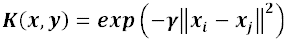 and sigmoidal: 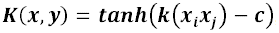, where *d* is the degree, **γ** > 0 sometimes parametrized as **γ** = **l**⁄**2*σ*^2^**, and *c* is a constant.

*Classification and regression trees:* Binary tree classifiers are built by repeated data splitting into two descendant subsets. Each terminal subset (node) is assigned a class label and the resulting partition of the data set corresponds to the classifier. There are three principal aspects of the tree building: (i) the selection of splitting rule; (ii) the split-stopping rule to declare a node terminal or continue splitting; (iii) the assignment of each terminal node to a class.

CART, which is introduced by Breiman [42], is one of the most popular tree classifiers and applied in many fields. It uses Gini index to choose the split that maximizes the decrease in impurity at each node. If *p(i|j)* is the probability of class *i* at node *j*, then the Gini index is 1 - ∑_i_ *p^2^(i|j)*. When CART grows a maximal tree, this tree is pruned upward to get a decreasing sequence of subtrees. Then, a cross-validation is used to identify the subtree that having the lowest estimated misclassification rate. Finally, the assignment of each terminal node to a class is performed by choosing the class that minimizes the resubstitution estimate of the misclassification probability [42, 43].

*Random forest:* A random forest is a collection of many CART trees combined by averaging the predictions of individual trees in the forest [44]. The idea behind the RF is to combine many weak classifiers to produce a significantly better strong classifier. For each tree, a training set is generated by bootstrap sample from the original data. This bootstrap sample includes 2/3 of the original data. The remaining of the cases are used as a test set to predict out-of-bag error of classification. If there are *m* features, *m_try_* out of m features are randomly selected at each node and the best split is used to split the node. Different splitting criteria can be used such as Gini index, information gain and node impurity. The value of *m_try_* is chosen to be approximately either 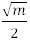 or 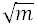 or 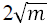 and constant during the forest growing. An unpruned tree is grown for each of the bootstrap sample, unlike CART. Finally, new data is predicted by aggregating, i.e. majority votes, the predictions of all trees [45, 46].

*Poisson linear discriminant analysis:* Let *X* be an *nxp* matrix of sequencing data, where *n* is number of observations and *p* is number of features. For sequencing data, *X_ij_* indicates the total number of reads mapping to gene *j* in observation *i*. Therefore, Poisson log linear model can be used for sequencing data,

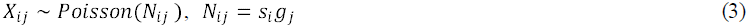

where *s_i_* is total number of reads per sample and *g_j_* is total number of reads per region of interest. For RNA-seq data, equation (3) can be extended as follows,

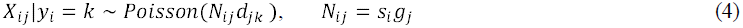

where 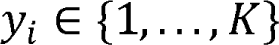 is the class of the *i^th^* observation, and *d_lj_,…,d_Kj_* terms allow the *j_th_* feature to be differentially expressed between classes.

Let (*x*_*i*_, *y*_*i*_), *i* = 1,…, *n*, be a training set and 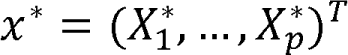 be a test set. Using the Bayes’ rule as follows,

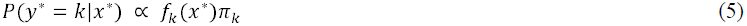

where *y*\* denotes the unknown class label, *f_k_* is the density of an observation in class *k* and *π_k_* is the prior probability that an observation belongs to class *k*. If *f_k_* is a normal density with a class-specific mean and common variance then a standard LDA is used for assigning a new observation to the class [47]. In case of the observations are normally distributed with a class-specific mean and a common diagonal matrix, then diagonal LDA methodology is used for the classification [48]. However, neither normality nor common covariance matrix assumptions are not appropriate for sequencing data. Instead, Witten [6] assumes that the data arise from following Poisson model,

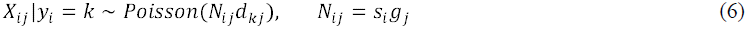

where *y_i_* represents the class of the *i^th^* observation and the features are independent.

The equation (4) specifies that 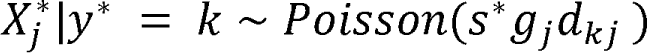. First, the size factors for the training data, *s*_1_,…, *s*_*n*_, is estimated. Then *s*\*, *g_j_*, *d_kj_* and *π_k_* are estimated as described in [6]. Substituting these estimations into equation (4) and recalling independent features assumption, equation (5) produces,

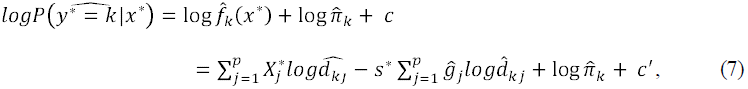

here *c* and *c*′ are constants and do not depend on the class label. The classification rule that assigns a new observation to the one of the classes for which equation (7) is the largest and it is linear in *x*\*. More detailed information can be found in [6].

### RNA-Seq Classification Workflow

Providing a pipeline for classification algorithm of RNA-Seq data gives us a quick snapshot view of how to handle the large-scale transcriptome data and establish a robust inference by using computer-assisted learning algorithms. Therefore, we outlined the count-based classification pipeline for RNA-Seq data in Figure 6. NGS platforms produce millions of raw sequence reads with quality scores corresponding to each base-call. The first step in RNA-Seq data analysis is to assess the quality of the raw sequencing data for meaningful downstream analysis. The conversion of raw sequence data into ready-to-use clean sequence reads needs a number of processes such as removing the poor-quality sequences, low-quality reads with more than five unknown bases, and trimming the sequencing adaptors and primers. In quality assessment and filtering, the current popular tools are FASTQC (http://www.bioinformatics.babraham.ac.uk/projects/fastqc/), HTSeq (http://www-huber.embl.de/users/anders/HTSeq/doc/overview.html), R ShortRead package [49], PRINSEQ (http://edwards.sdsu.edu/cgi-bin/prinseq/prinseq.cgi), FASTX Toolkit (http://hannonlab.cshl.edu/fastx_toolkit/) and QTrim [50]. Following these procedures, next step is to align the high-quality reads to a reference genome or transcriptome. It has been reported that the number of reads mapped to the reference genome is linearly related to the transcript abundance. Thus, transcript quantification (calculated from the total number of mapped reads) is a prerequisite for further analysis. Splice-aware short read aligners such as Tophat2 [51], MapSplice [52] or Star [53] can be prefered instead of unspliced aligners (BWA, Bowtie, etc.). After obtaining the mapped reads, next step is counting how many reads mapped to each transcript. In this way, gene expression levels can be inferred for each sample for downstream analysis. This step can be accomplished with HTSeq and bedtools [54] softwares. However, these counts cannot be directly used for further analysis and should be normalized to adjust between-sample differences. Moreover, for some applications such as differential expression, clustering and classification, a good way is to work with the transformed count data. Logarithmic transformation is the widely used choice; however it is probable to get zero count values for some genes. There is no standard tool for normalization, but the popular ones include deseq [17], trimmed mean of M values (TMM) [55], reads per kilobase per million mapped reads (RPKM) [56] and quantile [57]. For transformation, vst [17], rlog [9] and voom [31] methods can be a good choice. Once all mapped reads per transcripts are counted and normalized, we obtain gene-expression levels for each sample.

**Figure 6.**
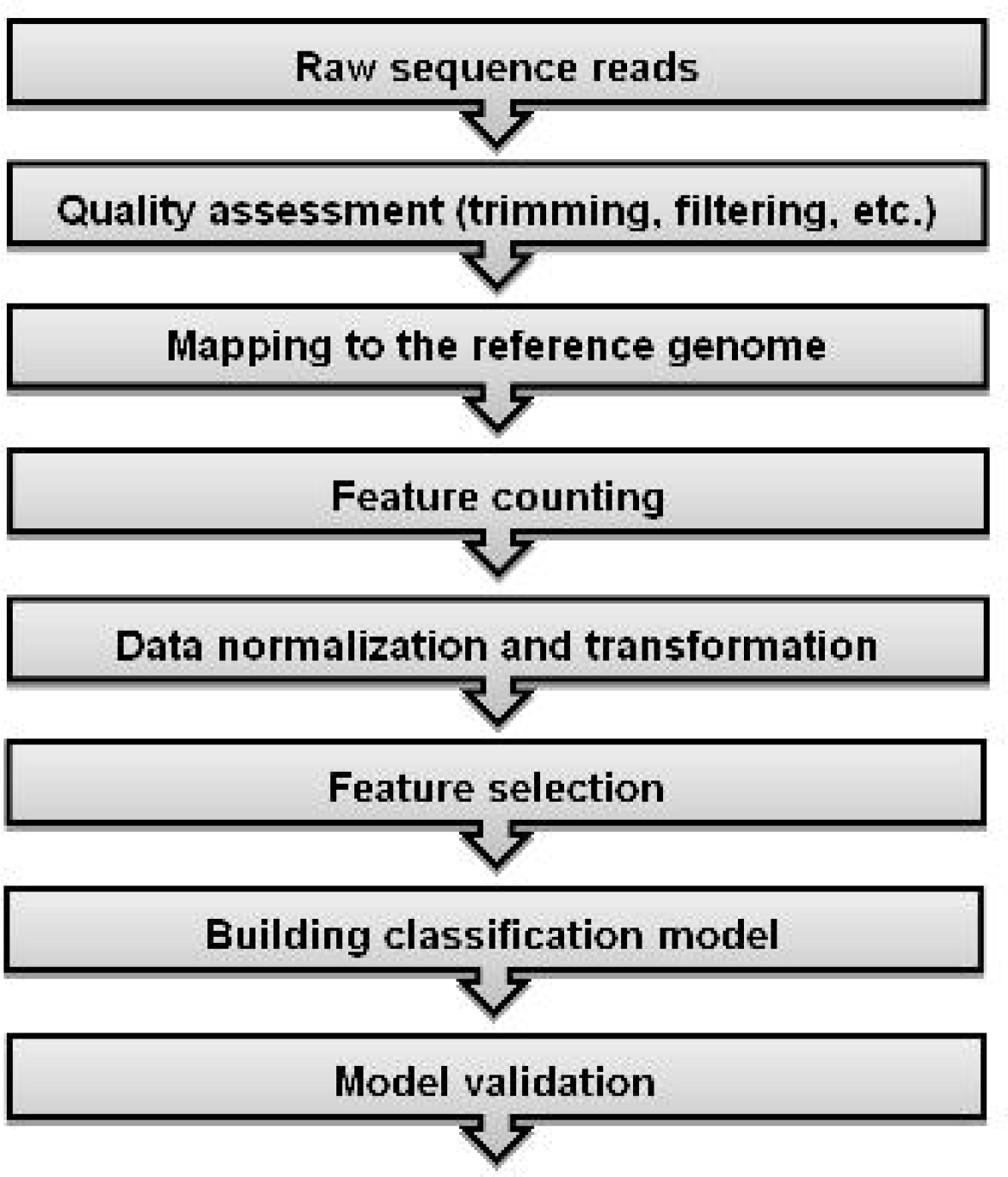
RNA-Seq classification workflow

The same workflow of microarray classification can be used after obtaining the gene-expression matrix. The crucial steps of classification can be written as feature selection, building classification model and model validation. In feature selection step, we aim to work with an optimal subset of data. This process is crucial to reduce the computational cost, decrease of noise and improve the accuracy for classification of phenotypes, also to work with more interpretable features to better understand the domain [58]. Various feature selection methods have been reviewed in detail and compared in [59]. Next step is model building, which refers to the application of a machine-learning algorithm and to learn the parameters of classifiers from training data. Thus, the built model can be used to predict class memberships of new biological samples. The commonly used classifiers include LDA, SVM, RF and other tree-based classifiers, artificial neural networks and k-nearest neighbors. In many real life problems, it is possible to experience that a classification algorithm may perform well and perfectly classify training samples, however perform poorly when classifying new samples. This problem is called as overfitting and independent test samples should be used to avoid overfitting and generalize classification results. Holdout, k-fold cross-validation, leave-one-out cross-validation and bootsrapping are among the recommended approaches for model validation.

## Authors’ contributions

GZ developed the method’s framework, DG and SK contributed to algorithm design and implementation. GZ, VE and IPD surveyed the literature for other available methods and collected performance data for the other methods used in the study for comparison. GZ, VE, IPD, DG carried out the simulation studies and data analysis. GZ, DG and SK developed MLSeq package. GZ, DG, SK and VE wrote the paper, TU and AO supervised the research process, revised and directed the manuscript. All authors read and approved the final manuscript.

## Acknowledgments

This work was supported by the Research Fund of Marmara University [FEN-C-DRP-120613-0273]. The authors declare that they have no competing interests.

